# Pro-cognitive Effects of Dual Tacrine Derivatives Acting as Cholinesterase Inhibitors and NMDA Receptor Antagonists

**DOI:** 10.1101/2024.03.18.585591

**Authors:** Marketa Chvojkova, David Kolar, Katarina Kovacova, Lada Cejkova, Anna Misiachna, Kristina Hakenova, Lukas Gorecki, Martin Horak, Jan Korabecny, Ondrej Soukup, Karel Vales

## Abstract

Therapeutic options for Alzheimer’s disease are limited. Dual compounds targeting two pathophysiological pathways concurrently may enable enhanced effect. The study focuses on tacrine derivatives acting as acetylcholinesterase (AChE) inhibitors and simultaneously as subunit-dependent N-methyl-D-aspartate (NMDA) receptor antagonists. Compounds with balanced inhibitory potencies for target proteins (K1578 and K1599) or with increased inhibitory potency for AChE (K1592 and K1594) were studied. We aimed to identify the most promising pro-cognitive compound.

The pro-cognitive effects of the compounds were studied in cholinergic (scopolamine-induced) and glutamatergic (MK-801-induced) rat models of cognitive deficits in the Morris water maze. Moreover, the effect on locomotion in open field and on AChE activity in relevant brain structures were investigated. The effect of the most promising compound on NMDA receptors was explored by *in vitro* electrophysiology.

The cholinergic antagonist scopolamine induced a deficit of memory acquisition, however was unaffected by the compounds, and a deficit of reversal learning, that was alleviated by K1578 and K1599. K1578 and K1599 significantly inhibited AChE in striatum, potentially explaining the behavioral observations.

Glutamatergic antagonist dizocilpine (MK-801) induced a deficit of memory acquisition, which was alleviated by K1599. K1599 also mitigated the MK-801-induced hyperlocomotion in the open field. The electrophysiology study corroborated the K1599-associated NMDA receptor inhibitory effect.

K1599 emerged as the most promising compound, demonstrating pro-cognitive efficacy in both models, consistently with intended dual effect. Our findings contributed to elucidation of structural and functional properties of tacrine derivatives associated with optimal *in vivo* pro-cognitive effects, which further research may benefit from.

## 1 Introduction

Alzheimer’s disease (AD), a prevalent neurodegenerative disorder causing dementia ^1^, poses a growing socio-economic challenge due to population aging and limited therapeutic success ^2^. This multifactorial disease involves a complex pathophysiology ^2^, including cholinergic neuron loss and, apparently, N-methyl-D-aspartate (NMDA) glutamate receptor overstimulation ^3–6^. However, the pathophysiological processes are widely interconnected ^6–9^ and together drive disease progression.

Current therapy offers only palliative effects via acetylcholinesterase (AChE) inhibitors, indirectly stimulating the cholinergic system, and the NMDA receptor antagonist memantine, assumed to mitigate the NMDA receptor overactivation ^10^. From other experimental approaches, the single-target strategies have not succeeded ^2^ except questionable recently presented immunotherapies ^11^. Given the disease’s complexity, a polypharmacology approach, targeting multiple pathways concurrently, emerges as a more promising strategy ^12,13^. This strategy aims to enhance therapeutic efficacy while minimizing side effects ^10,12,14,15^, particularly through simultaneous AChE and NMDA receptor inhibition ^10,13,14^.

Co-administration of AChE inhibitors and NMDA receptor antagonists enhanced cognitive recovery in rodent models ^16,17^ as well as in patients ^15,18^, leading to drug approval (Namzaric) ^19^. This combination therapy sets the stage for the development of multi-target drugs, single drugs designed to act on multiple targets and overcome some limitations of combination therapy ^12–14^.

Tacrine, the first AChE inhibitor for AD, displayed therapeutic potential ^20^ but was discontinued due to hepatotoxicity risk ^21^. However, efforts to develop safer derivatives continue ^22,23^. Interestingly, tacrine also exhibits NMDA receptor inhibition ^24,25^. Its dual actions and low molecular weight make it a valuable starting point for development of dual-target derivatives with improved properties (**Figure 1a**). In particular 7-methoxytacrine (7-MEOTA; **Figure 1b**) or 7-phenoxytacrine represent promising lead dual derivatives with improved neuroprotective properties ^25,26^.

**Figure 1.**
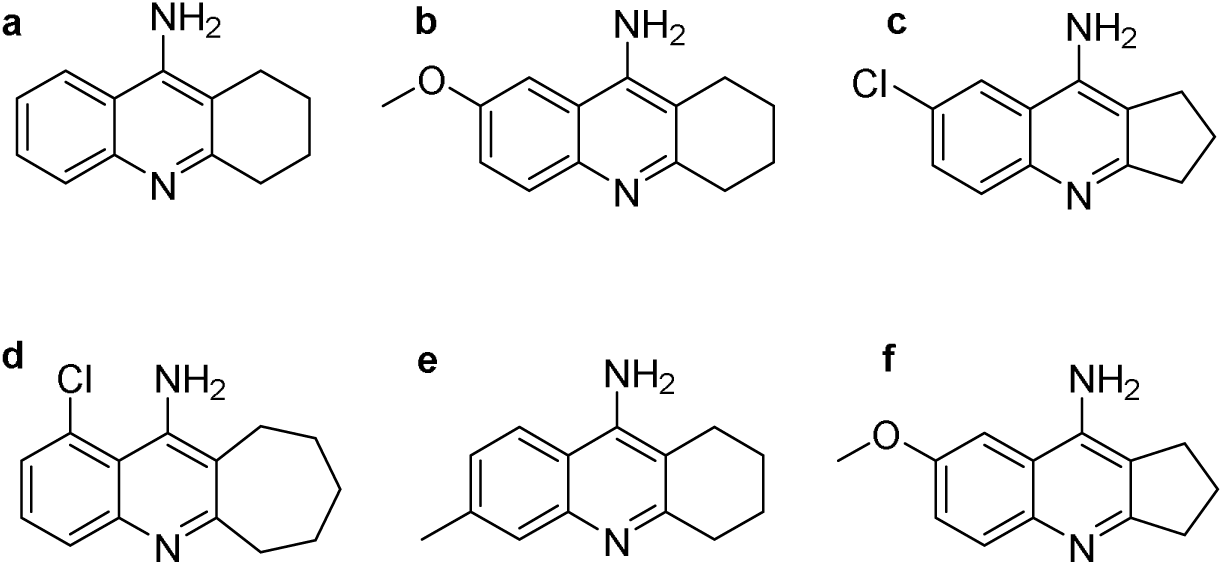
Chemical structures of tacrine (**a**) and its derivatives 7-MEOTA (**b**), K1578 (7-chloro-1*H*,2*H*,3*H*-cyclopenta[*b*]quinolin-9-amine; **c**), K1592 (1-chloro-6*H*,7*H*,8*H*,9*H*,10*H*-cyclohepta[*b*]quinolin-11-amine; **d**), K1594 (6-methyl-1,2,3,4-tetrahydroacridin-9-amine; **e**) and K1599 (7-methoxy-1*H*,2*H*,3*H*-cyclopenta[b]quinolin-9-amine; **f**). Compounds in this study were used in the form of hydrochloride salts.

In our previous work, we introduced tacrine derivatives with varying affinities for NMDA receptor subtypes (GluN1/GluN2A vs. GluN1/GluN2B) and cholinesterases (AChE and butyrylcholinesterase, BChE) ^27^ due to the fact that distinct NMDA receptor subtypes play different roles in pathology ^28,29^. Building on these findings, four compounds (K1578, K1592, K1594, and K1599) were selected for the current extensive *in vivo* efficacy study, emphasizing cognition restoration and potential dual effects (**Figure 1c–f**).

Generally, a balanced target affinity, specifically AChE and NMDA receptor inhibition, is a prerequisite for multi-target directed ligands ^14^. On the other hand, a tiny difference in the affinity as well as the subtype selectivity may result in a completely different phenotype *in vivo*. K1578 and K1599 demonstrated rather balanced inhibitory potencies towards NMDA receptors and cholinesterases, with the IC_50_ values in the one- to two-digit micromolar range. The IC_50_ values of K1578 followed the order AChE < BChE < GluN1/GluN2B < GluN1/GluN2A, while those of K1599 displayed the GluN1/GluN2A < AChE < BChE < GluN1/GluN2B relationship. K1592 and K1594 also were micromolar NMDA receptor inhibitors, but with a strong, submicromolar, inhibitory potency for AChE, even surpassing that of tacrine. K1592 was also a submicromolar inhibitor of BChE. Regarding NMDA receptors, K1592 and K1594 showed a slight preference for GluN1/GluN2A and GluN1/GluN2B, respectively (For exact IC_50_ values see ^27^.) Of note, all the selected compounds are able to cross the blood-brain barrier and are available in the brain *in vivo* ^27^.

The main aim of the current study was to investigate and compare their pro-cognitive effects, given by the cholinergic and glutamatergic characteristics, in cholinergic and glutamatergic rat models of cognitive deficit, i.e. induced by scopolamine, a competitive antagonist of muscarinic acetylcholine receptors ^30^ or dizocilpine (MK-801), a potent NMDA receptor antagonist ^31^, respectively. The experiments assessed spatial memory and cognitive flexibility, both relevant to AD ^32,33^, and included additional analyses such as open field behavior, patch-clamp study, and AChE inhibition in specific brain regions to elucidate the compoundś mechanisms of action. The pro-cognitive, behavioral, AChE- and NMDA receptor-inhibitory effects of the parent compound tacrine in aforementioned assays are known from literature ^25,34–36^. Finally, the most beneficial tacrine derivative effective in both models was selected.

## 2 Materials and methods

### 2.1 Animals

Adult male Wistar rats (*Rattus norvegicus;* 280–460 g, 2–3 months) from the Velaz breeding facility (Czech Republic) were used. They were housed in pairs in transparent plastic boxes (23 × 38 × 23 cm) in an animal room of National Institute of Mental Health, Czech Republic, with constant temperature (22.5 °C), humidity, and 12:12 h light/dark cycle, with free access to water and food pellets. Experiments were performed in the light phase of the day after a week-long acclimatization period. The experiments were conducted in accordance with the guidelines of European Union directive 2010/63/EU and Act No. 246/1992 Coll. on the protection of animals against cruelty, and were approved by the Animal Care and Use Committee of the National Institute of Mental Health (reference number MZDR 51755/2018-4/OVZ). In determining the sample size, relevant literature pertaining to corresponding methodologies ^37–39^ as well as the principles of Replacement, Reduction, and Refinement of animals used in research were considered.

### 2.2 Drugs and administration to animals

Compounds K1578, K1592, K1594 and K1599 (in the form of hydrochloride salts) synthesized according to ^27^ were used (HPLC purity > 97%). Other compounds for animal administration were purchased from Sigma-Aldrich (St. Louis, MO, USA). Doses are expressed as the salt form of the drugs.

#### Administration of compounds

Compounds K1578, K1592, K1594, and K1599 were administered at 1 mg/kg by dissolving them in 5% dimethyl sulfoxide (DMSO) in physiological saline, resulting in a 1 mg/mL concentration, with an injection volume of 1 mL/kg. For the 5 mg/kg dose, the compounds were dissolved in the same vehicle at a 2 mg/mL concentration, administered at 2.5 mL/kg. MK-801 ((+)-MK-801 hydrogen maleate; 0.2 mg/kg for the open field test, 0.1 mg/kg for Morris water maze; MWM) and scopolamine hydrobromide (2 mg/kg) were dissolved in physiological saline at 1 mL/kg. All compounds were administered intraperitoneally (ip); the route and doses were chosen based on previous experience ^27,40,41^. Co-administration of the 5 mg/kg dose of K1592 with MK-801 was not performed due to side effects.

#### Timing of administration

In the behavioral experiments, MK-801 was administered 30 min, scopolamine 20 min, and K1578, K1592, K1594, and K1599 15 min before the start of the behavioral testing, based on their pharmacokinetic properties ^27^. In the AChE enzyme activity assay, the drugs were administered 30 min before euthanizing the subjects, which reflects the timing during the MWM testing.

### 2.3 Behavioral experiments

#### 2.3.1 Morris water maze

##### Treatment groups

The rats were pseudo-randomly assigned to one of the 18 treatment groups listed in **Table 1** (upper part). Each group received two injections: one containing the study compound and another containing either MK-801 or scopolamine, as indicated by the group name. The vehicle group (VEH) received the DMSO vehicle (2.5 mL/kg) and saline. The “scopolamine” and “MK-801” groups received scopolamine or MK-801, respectively, along with the DMSO vehicle (2.5 mL/kg).

**Table 1:** Treatment groups and *n* in behavioral experiments.

| Groups | *n* | Groups | *n* |
| --- | --- | --- | --- |
| **Morris water maze** |  |  |  |
| **Scopolamine model** |  | **MK-801 model** |  |
| VEH (shared for both models) | 7 |  |  |
| Scopolamine | 7 | MK - 801 | 6 |
| K1578 (1 mg/kg) + scopolamine | 6 | K1578 (1 mg/kg) + MK-801 | 6 |
| K1578 (5 mg/kg) + scopolamine | 6 | K1578 (5 mg/kg) + MK-801 | 6 |
| K1592 (1 mg/kg) + scopolamine | 6 | K1592 (1 mg/kg) + MK-801 | 6 |
| K1592 (5 mg/kg) + scopolamine | 6 |  |  |
| K1594 (1 mg/kg) + scopolamine | 6 | K1594 (1 mg/kg) + MK-801 | 5 |
| K1594 (5 mg/kg) + scopolamine | 6 | K1594 (5 mg/kg) + MK-801 | 6 |
| K1599 (1 mg/kg) + scopolamine | 9 | K1599 (1 mg/kg) + MK-801 | 6 |
| K1599 (5 mg/kg) + scopolamine | 9 | K1599 (5 mg/kg) + MK-801 | 6 |
| **Open field** |  |  |  |
| **Intact phenotype** |  | **MK-801 phenotype** |  |
| VEH (1 ml/kg) | 8 | VEH (1 ml/kg) + MK-801 | 6 |
| VEH (2.5 ml/kg) | 6 | VEH (2.5 ml/kg) + MK-801 | 7 |
| K1578 (1 mg/kg) | 6 | K1578 (1 mg/kg) + MK-801 | 6 |
| K1578 (5 mg/kg) | 6 | K1578 (5 mg/kg) + MK-801 | 8 |
| K1592 (1 mg/kg) | 6 | K1592 (1 mg/kg) + MK-801 | 6 |
| K1594 (1 mg/kg) | 6 | K1594 (1 mg/kg) + MK-801 | 6 |
| K1594 (5 mg/kg) | 6 | K1594 (5 mg/kg) + MK-801 | 6 |
| K1599 (1 mg/kg) | 6 | K1599 (1 mg/kg) + MK-801 | 6 |
| K1599 (5 mg/kg) | 6 | K1599 (5 mg/kg) + MK-801 | 6 |

Note: *n*: number of subjects per group.

##### Morris water maze apparatus

MWM is a routinely used test of spatial cognition for rodents ^42^. The apparatus consisted of a 180 cm diameter gray plastic pool with a hidden platform (circular, 10 cm diameter, transparent, submerged 1 cm below the water surface). The water (40 cm depth) was maintained at 23°C and tinted with non-toxic gray dye. No visual cues were present on the maze walls, requiring rats to rely on constant room-based cues for navigation.

##### Testing procedure

The experiment spanned 5 consecutive days. The drugs were administered daily before testing. On d 1–4, the position of the hidden platform remained constant in the center of a selected quadrant for acquisition trials. On d 5, the hidden platform was moved to the center of the opposite quadrant for reversal trials ^42,43^. Reversal learning reflects cognitive flexibility, a complex process enabling adjustment of subject’s behavior to the changes of the environment (switching the search strategy to the new platform position) ^42,44,45^. Each day, rats underwent 8 swims from different starting positions at the perimeter of the pool (labeled N, S, E, W, NE, NW, SE, SW, pseudo-random order) with an intertrial interval of 4–7 min. Rats were placed into the pool facing its inner wall and given 60 s to locate the platform. If a rat failed to find the platform, it was gently guided to it by the experimenter. The rat remained on the platform for 15 s to acquire spatial cues before being removed. No evident motor impairment was observed in the rats during the MWM testing.

##### Data collection

Rat positions were monitored using a camera mounted above the pool, connected to a tracking software (EthoVision 11, Noldus, Netherlands). The dependent variable was escape latency (the latency to find the platform), represented as the daily mean value. In cases where a rat did not find the platform, a latency of 60 s was recorded.

#### 2.3.2 Open field test

##### Treatment groups

The rats were pseudo-randomly assigned to one of the 18 treatment groups listed in **Table 1** (lower part). Animals received 1 (intact phenotype groups) or 2 (MK-801 phenotype groups) injections, as indicated by their group names. (Due to side effects, co-administration of MK-801 with the 5 mg/kg dose of K1592 was not conducted.) The “VEH” groups received the specified volumes of the DMSO vehicle. The “VEH + MK-801” groups received the DMSO vehicle along with MK-801. Two different volumes of the DMSO vehicle were used in the control animals, to provide appropriate control groups for the 1 and 5 mg/kg treatment groups. However, as there was not statistically significant difference between the distance moved by the group “VEH 1 mL/kg” and “VEH 2.5 mL/kg”, as well as between “VEH 1 mL/kg + MK-801” and “VEH 2.5 mL/kg + MK-801” (t-test), these groups were merged and referred to as “VEH” and “VEH + MK-801”, respectively.

##### Data source clarification

Data from the intact phenotype groups (i.e. without MK-801 co-administration) have been previously published for side effect assessment in ^27^. These experiments were not replicated in the current publication to minimize the use of laboratory animals.

##### Open field test apparatus and procedure

The open field test used a black plastic square arena (80 × 80 cm), located in a controlled-light room. Rats were placed in the center of the arena, and their activity was tracked for 10 min using a camera connected to tracking software (EthoVision 14, Noldus, Netherlands). The arena was thoroughly cleaned after each animal. The dependent variable was the distance moved by the animal.

### 2.4 Biochemistry experiments - *In situ* acetylcholinesterase activity assay

#### 2.4.1 Sacrifice and tissue preparation

Separate sets of experimentally naïve rats were administered with the compounds at 1 mg/kg via the same route and at a corresponding time point as the rats subjected to MWM. The rats were pseudo-randomly assigned in the groups listed in **Table 2**. The VEH groups, receiving the DMSO vehicle in the volume of 1 mL/kg or 2.5 mL/kg, were merged into one control group as no statistically significant alterations of AChE activity within the analyzed brain regions were observed between the groups (data not shown).

**Table 2:** Treatment groups and *n* in biochemical experiments - AChE activity assay.

| Groups | *n* |
| --- | --- |
| VEH (1 ml/kg) | 6 |
| VEH (2.5 ml/kg) | 8 |
| K1578 (1 mg/kg) | 10 |
| K1592 (1 mg/kg) | 6 |
| K1594 (1 mg/kg) | 6 |
| K1599 (1 mg/kg) | 10 |

Note: *n*: number of subjects per group.

All rats were euthanized via decapitation in accordance with ethical protocols. Thirty minutes after drug administration, the decapitation procedure was conducted humanely, and the brains were swiftly excised and rinsed with ice-cold saline. The brain was divided sagittally into two halves. The prefrontal cortex, striatum, and hippocampus, structures involved in the studied cognitive functions ^45–47^, were dissected from the left half of the brain, while the entire right half of the brain (including the right half of the cerebellum) was left intact and referred to as whole brain homogenate.

The collected tissue samples were weighed, immediately frozen at dry ice and stored at –80 °C. The samples were homogenized on ice solubilization solution composed of 10 mM TRIS, 1 M NaCl, 50 mM MgCl_2_, 1% Triton X-100, and a cocktail of protease (cOmplete®, Roche) and phosphatase (PhosSTOP®, Roche) inhibitors (pH = 7.2). This homogenization process employed an IKA® ULTRA-TURRAX® with dispersing elements S10N-5G and 8-G, resulting in 10% (w/v) tissue homogenates. Right upon collecting the homogenate, a small portion of the homogenate was used to determine the protein concentration using the Bradford dye (Merck) binding assay (Bradford, 1976). The remaining homogenate was appropriately labeled, frozen and stored for subsequent assessment of AChE enzyme activity.

#### 2.4.2 Acetylcholinesterase enzyme activity assessment

Despite the concurrent BChE-inhibiting effects of the compounds, we focused an AChE activity only, as BChE expression in the key structures, hippocampus and striatum, is marginal or absent ^48,49^. The enzymatic activity of AChE was assessed using the spectrophotometric Ellman method, a widely recognized approach for quantifying cholinesterase activity ^50^. This method relies on the reaction between acetylthiocholine iodide and 5,5′-dithiobis-2-nitrobenzoic acid (DTNB), generating a yellow product comprising mercapto-2-nitrobenzoic acid and its dissociated forms. This chemical transformation occurs under controlled conditions at pH 8.0, with the highest absorption coefficient observed at 412 nm, having a specific value of 13.6 × 10^−3^ M^−1^ cm^−1^. Briefly, thawed sample aliquots were significantly diluted (10–15 times) with a solubilization solution, resulting in a final volume of 100 μL per sample. These diluted samples were then dispensed in triplicate into a 96-well microplate. To provide a basis for comparison, two control groups were included: one contained 100 μL of the solubilization solution (the negative control), and the other contained 100 μL of diluted AChE enzyme sourced from Electrophorus electricus (Merck) as the positive control. Following this, 50 μL of a 1.25 mM DTNB solution, prepared in 0.1 M Phosphate buffer at pH 8.0, was added to each well. Subsequently, the samples were incubated for 5 min at room temperature. Next, the microplate was transferred to a microplate reader, specifically the Infinite M200Pro by Tekan®. The enzymatic reaction was initiated by introducing 50 μL of an acetylthiocholine iodide solution, prepared at a concentration of 1.75 mM in 0.1 M Phosphate buffer with a pH of 8.0. The reaction kinetics, involving the increase in absorbance at λ = 412 nm, were monitored and recorded over a period of 10 min, with readings taken at 1-min intervals. Subsequently, the specific activity of the samples was computed as activity units per milligram of protein present in each well. Finally, the enzyme activities within the tacrine derivative groups were expressed relative to the control group, quantified as fold changes in comparison to the control group.

### 2.5 *In vitro* electrophysiology experiments

Whole-cell patch-clamp measurements were performed on transfected HEK293 cells expressing rat version of the GluN1-4a/GluN2A (GluN1/GluN2A) receptors using an Axopatch 200B amplifier (Molecular Devices) as described ^25,51^. The cells were placed in an extracellular solution (ECS) containing (in mM): 160 NaCl, 2.5 KCl, 10 HEPES, 10 D-glucose, 0.2 EDTA, and 0.7 CaCl_2_ (pH 7.3 with NaOH). Glass patching pipettes (3–6 MΩ resistance) were prepared using a P-1000 puller (Sutter Instruments) and filled with intracellular solution (ICS) containing (in mM): 125 gluconic acid, 15 CsCl, 5 EGTA, 10 HEPES, 3 MgCl_2_, 0.5 CaCl_2_, and 2 ATP-Mg salt (pH 7.2 with CsOH). Electrophysiological recordings were conducted at room temperature using membrane potentials from −80 to 60 mV (with a step of 20 mV); the junction potential was not subtracted. The GluN1/GluN2A receptor currents were induced by fast application of 1 mM glutamate in the continuous presence of 100 μM glycine. The stock solution of K1599 was freshly dissolved in DMSO and diluted to the final working concentration of 30 μM. All chemicals described above were obtained from Merck. Data were analyzed using Clampfit 10.2 software (Molecular Devices).

### 2.6 Statistics

Data from individual experiments were examined separately due to variations in the protocols employed. The data from MWM reversal and AChE activity assay were analyzed using one-way analysis of variance (ANOVA) followed by Dunnett‘s multiple comparisons test when appropriate. The data from the open field were analyzed using two-way ANOVA (factors: tacrine derivative treatment and phenotype i.e. MK-801) with Dunnett‘s multiple comparisons test (to investigate treatment effect; comparison vs. VEH group of the corresponding phenotype). Aforementioned data underwent rigorous scrutiny, involving checks for outliers using the robust regression and outlier removal (ROUT) method with a Q value set at 1%, and assessments for normality (Shapiro-Wilk test, Kolmogorov-Smirnov test) and for homogeneity of variances (Brown-Forsythe test). In cases where the assumption of homogeneity of variances was not met, Welch’s ANOVA was employed, followed by corrections for multiple comparisons using Dunnett’s T3 multiple comparisons test. The mean daily latencies from MWM acquisition were analyzed using two-way repeated measures ANOVA (day as repeated factor) followed by Dunnett‘s multiple comparisons test (to investigate main treatment effect by comparing mean latencies over all acquisition days). In behavioral experiments, separate ANOVAs were performed by compounds. The predetermined level of significance was established at p ≤ 0.05. All statistical analyses were executed using GraphPad Prism 8 software (San Diego, USA).

## 3 Results

### 3.1 Morris water maze

#### 3.1.1 Scopolamine-induced model of cognitive deficit

##### 3.1.1.1 Acquisition

Two-way repeated measures ANOVAs of the escape latencies on the MWM acquisition days were conducted separately for the K1578, K1592, K1594, and K1599 groups, along with their respective control groups. These analyses revealed significant effects of treatment (F (3, 22) = 8.737, p = 0.0005; F (3, 22) = 8.801, p = 0.0005; F (3, 22) = 6.026, p = 0.0037; and F (3, 28) = 5.424, p = 0.0045, respectively) and testing day (F (3, 66) = 29.18; F (3, 66) = 30.19; F (3, 66) = 38.86; and F (3, 84) = 48.91, respectively; p < 0.0001 for all). We observed decreasing escape latencies along with the increase of testing day as the rats learned. No significant interactions between treatment and day factors were observed.

Post hoc tests were focused on the main treatment effect, which involved examining mean group latencies over all the acquisition days. These tests revealed a significantly increased mean latency in the scopolamine group compared to the VEH group (p < 0.01), indicating a deficit in spatial learning.

The groups co-administered with scopolamine and K1578 (p = 0.0045 for 1 mg/kg dose and p = 0.0004 for 5 mg/kg; **Figure 2a**), K1592 (p = 0.0102 for 1 mg/kg and p = 0.0002 for 5 mg/kg; **Figure 2b**), K1594 (p = 0.0074 for 1 mg/kg and p = 0.0068 for 5 mg/kg; **Figure 2c**) or K1599 (p = 0.0163 for 1 mg/kg and p = 0.0037 for 5 mg/kg; **Figure 2d**) displayed increased latencies compared to VEH. This indicates that all the compounds failed to reduce the scopolamine-induced deficit of spatial learning.

**Figure 2.**
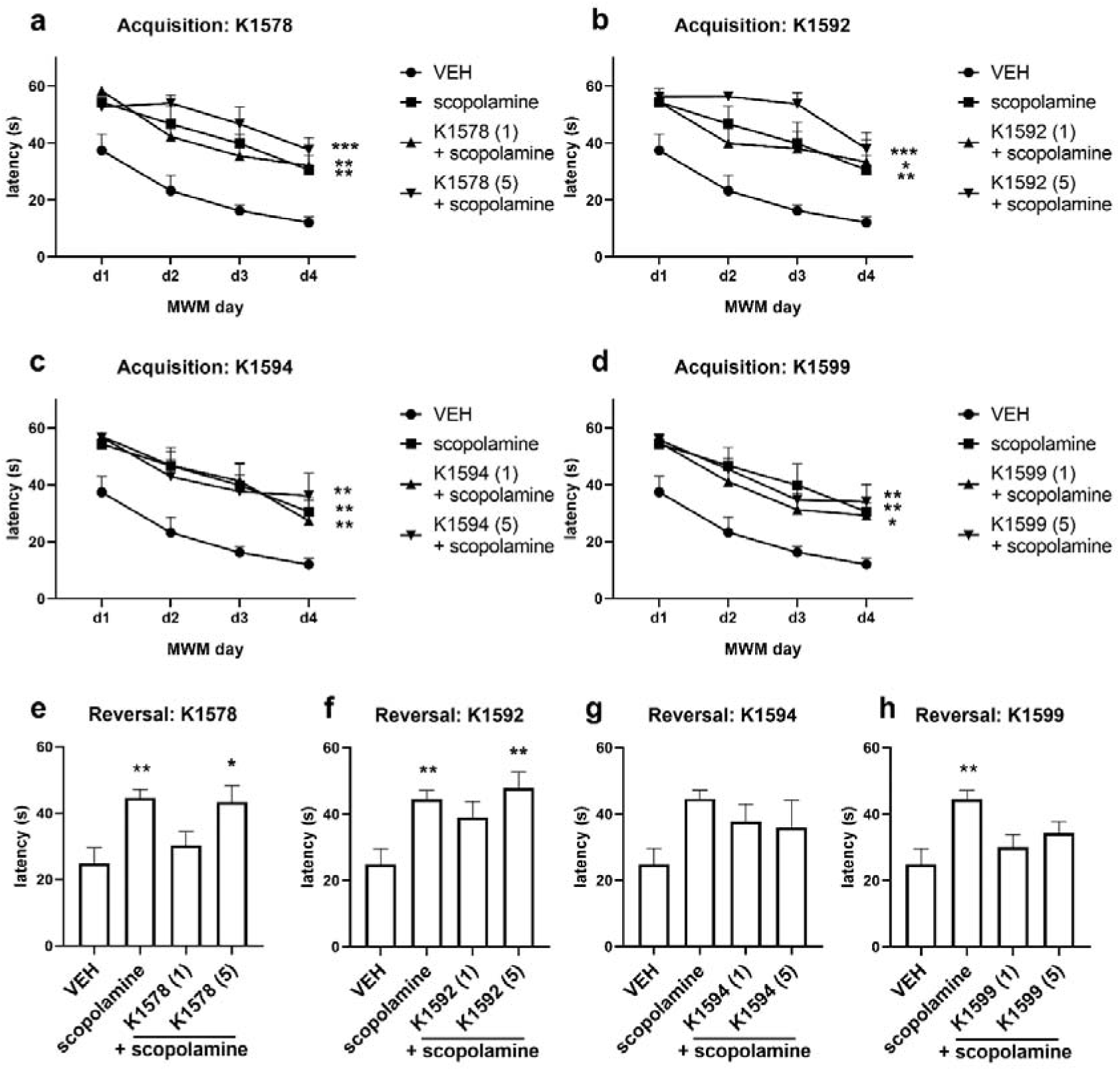
Morris water maze: scopolamine-induced model of cognitive deficit in the acquisition and reversal phases. The graphs show the effects of K1578 (**a**), K1592 (**b**), K1594 (**c**) and K1599 (**d**) on escape latency in the acquisition phase, where none of the compounds ameliorated the deficit of spatial learning. Remaining graphs display the effects of K1578 (**e**), K1592 (**f**), K1594 (**g**) and K1599 (**h**) in the reversal, where K1578 (1 mg/kg) and K1599 (both doses), and marginally also K1592 (1 mg/kg; see in the text) mitigated the scopolamine-induced deficit of reversal learning. VEH – vehicle, the numbers in the brackets denote the dose applied (mg/kg). Data are represented as the mean + SEM, * vs. VEH, * p < 0.05, ** p < 0.01, *** p < 0.001. *n* = 6–9 animals per group.

##### 3.1.1.2 Reversal

On the 5th day the platform was moved to the opposite quadrant to study reversal learning. ANOVAs of the mean escape latencies of the K1578 groups (F (3, 22) = 5.650, p = 0.0050), K1592 groups (F (3, 22) = 5.989, p = 0.0038), and K1599 groups (F (3, 28) = 4.631, p = 0.0094) revealed a treatment effect. The post hoc tests showed that scopolamine increased the escape latency (p < 0.01 vs. VEH), indicating a deficit of reversal learning.

Regarding K1578 groups, the post hoc test showed that the latency of the K1578 (1 mg/kg) + scopolamine group did not significantly differ from VEH (p = 0.6901). It suggests mitigation of the scopolamine-induced cognitive deficit by K1578. However, the animals treated with the higher dose of K1578 (5 mg/kg) showed increased latencies (p = 0.0130 vs. VEH), indicating that this dose of K1578 did not mitigate the cognitive deficit (**Figure 2e**).

Similarly, the post hoc test focused on the K1592 groups revealed that the latency of the K1592-treated animals (1 mg/kg) did not differ from VEH, suggesting possible mitigation of cognitive deficit. However, the reduction of the cognitive deficit was only minor, as seen from the group means (**Figure 2f**). Considering the 5 mg/kg dose of K1592, this group again showed an increased latency (p = 0.0025 vs. VEH), indicating no mitigation of cognitive deficit (**Figure 2f**).

In K1599 groups, the post hoc test showed that the latency of the group K1599 (1 mg/kg) + scopolamine as well as K1599 (5 mg/kg) + scopolamine did not differ from VEH (p = 0.6329 and 0.1881, respectively), suggesting alleviation of cognitive deficit by both doses of K1599 (**Figure 2h**).

In the case of K1594 groups, ANOVA found no differences between the groups, therefore the post hoc test could not be performed (**Figure 2g**).

To sum up, K1578 (1 mg/kg) and K1599 (both doses), and marginally also K1592 (1 mg/kg) mitigated the scopolamine-induced deficit of reversal learning.

#### 3.1.2 MK-801-induced model of cognitive deficit

##### 3.1.2.1. Acquisition

Separate two-way repeated measures ANOVAs of the escape latencies of the K1578, K1592, K1594 and K1599 groups revealed the effect of treatment (F (3, 21) = 8.155, p = 0.0009; F (2, 16) = 9.957, p = 0.0016; F (3, 20) = 6.459, p = 0.0031; and F (3, 21) = 5.263, p = 0.0073, respectively) and testing day (F (3, 63) = 31.07; F (3, 48) = 36.15, F (3, 60) = 26.17; and F (3, 63) = 40.36, respectively; p < 0.0001 for all), with no interaction between these factors. The post hoc tests focused on the main treatment effect showed increased mean latency of the MK-801 group (p < 0.01 vs. VEH), clearly confirming a deficit of spatial learning.

The groups co-administered with MK-801 and K1578 (p = 0.0013 for 1 mg/kg dose and p = 0.0039 for 5 mg/kg; **Figure 3a**), K1592 (1 mg/kg; p = 0.0047; **Figure 3b**), or K1594 (p = 0.0033 for 1 mg/kg and p = 0.0161 for 5 mg/kg; **Figure 3c**), respectively, displayed significantly increased latencies compared to VEH as well. It indicates that these compounds did not mitigate the MK-801-induced cognitive deficit.

**Figure 3.**
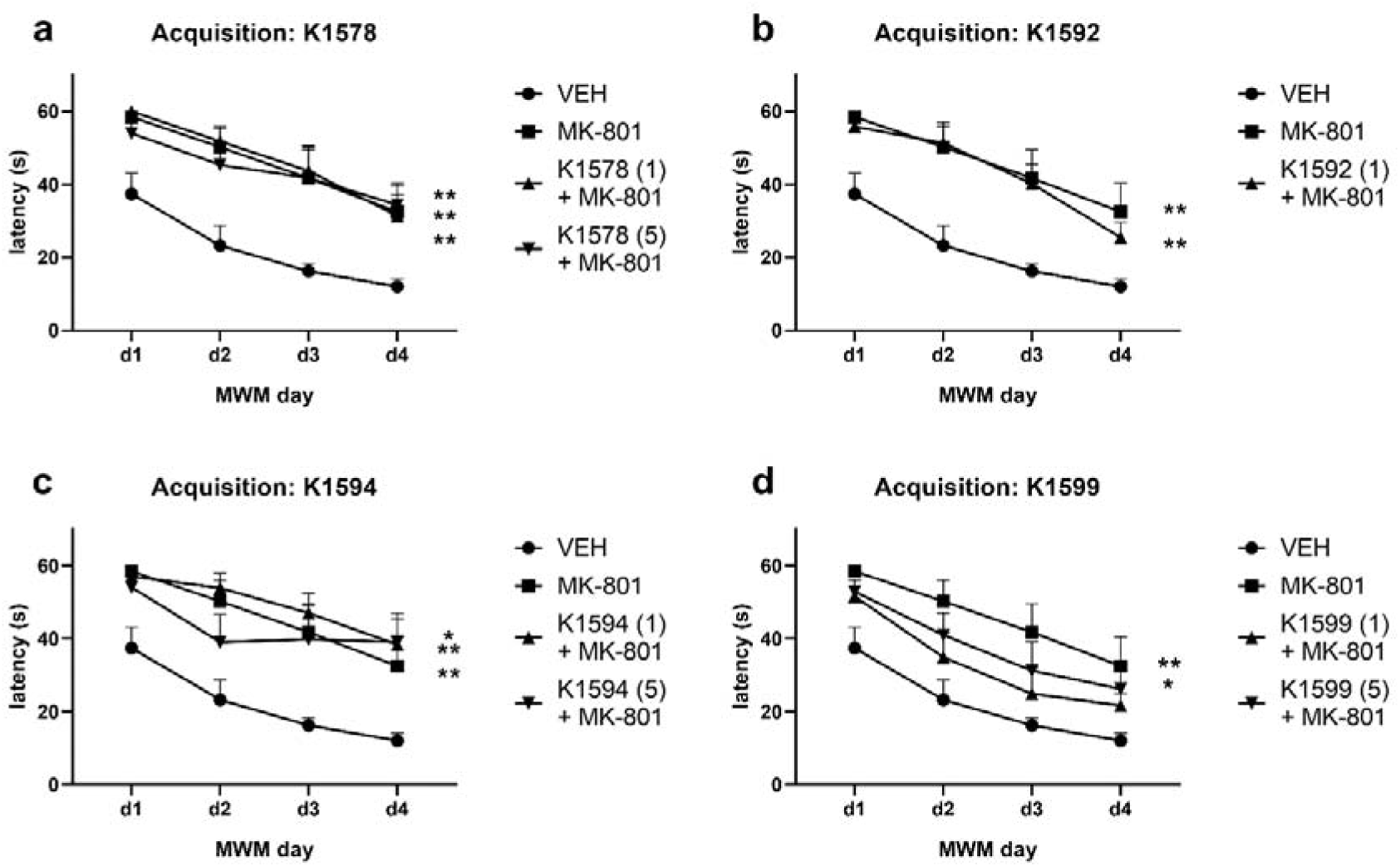
Morris water maze: MK-801-induced model of cognitive deficit in the acquisition phase. The graphs show the effects of the compounds K1578 (**a**), K1592 (**b**), K1594 (**c**) and K1599 (**d**) on escape latency. Only K1599 (1 mg/kg) ameliorated the MK-801-induced deficit of spatial learning. VEH – vehicle, the numbers in the brackets denote dose (mg/kg). Data are represented as the mean + SEM, * vs. VEH, * p < 0.05, ** p < 0.01. *n* = 5–7 animals per group.

However, animals co-treated with MK-801 and K1599 at the 1 mg/kg dose did not exhibit a significant difference in latency compared to VEH (p = 0.2017), suggesting an improvement of the MK-801-induced cognitive deficit by K1599 at this lower dose. Notably, at the 5 mg/kg dose of K1599, the animals showed increased latency (p = 0.0475 vs. VEH), indicating a lack of ameliorative effect of K1599 at this dose (**Figure 3d**).

In summary, only K1599 at the 1 mg/kg dose demonstrated an ameliorative effect on the MK-801-induced deficit of spatial learning.

##### 3.1.2.2 Reversal

In the reversal phase of the experiment (see above), MK-801 failed to induce a statistically significant cognitive deficit, therefore it was not possible to study the effects of the drugs in this part of the experiment (data not shown). ANOVAs of the latencies of the K1578, K1592, K1594, as well as K1599 groups failed to show significant differences among the means.

### 3.2 Open field test

To further investigate NMDA receptor-mediated actions, the effects of the compounds on locomotor activity were tested in two rat phenotypes: intact and MK-801-treated rats, where MK-801 induced hyperlocomotion.

Analysis of the distance moved by K1578 groups revealed the effect of K1578 (F (2, 46) = 75.11, p < 0.0001), MK-801 (F (1, 46) = 144.8, p < 0.0001), and interaction of these factors (F (2, 46) = 5.111, p = 0.0099). K1578 (1 mg/kg) did not affect locomotion of intact animals, but it further increased the hyperlocomotion in MK-801-treated animals (p = 0.0165 vs. VEH+MK-801). However, at 5 mg/kg, K1578 decreased locomotion of the intact as well as MK-801-treated rats (p < 0.0001 for both; **Figure 4a**).

**Figure 4.**
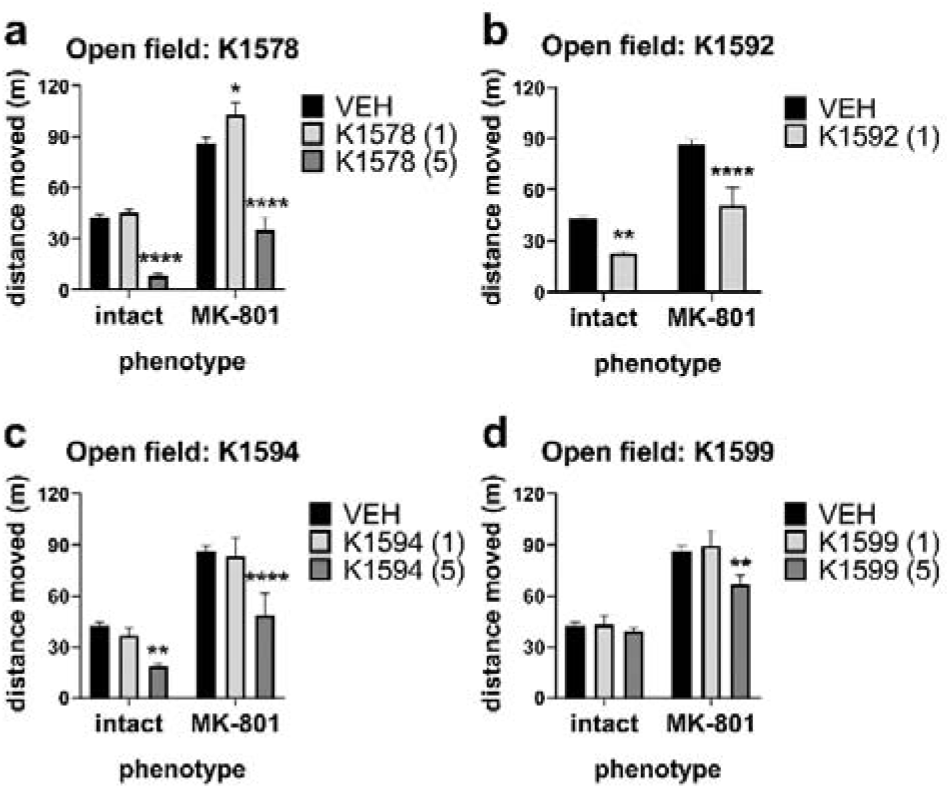
Open field test. The graphs show the effects of K1578 (**a**), K1592 (**b**), K1594 (**c**), and K1599 (**d**) on the distance moved by intact and MK-801-treated animals. VEH – vehicle, the numbers in the brackets denote dose (mg/kg). Data are represented as the mean + SEM, * vs. VEH group of the corresponding phenotype, * p < 0.05, ** p < 0.01, **** p < 0.0001. *n* = 6–14 animals per group.

Analysis of the K1592 groups yielded the effect of K1592 and of MK-801 (F (1, 34) = 40.19 and F (1, 34) = 67.16, respectively, p < 0.0001 for both). K1592 (1 mg/kg) decreased locomotion of the intact animals (p = 0.0049) and of the MK-801-treated animals (p < 0.0001; **Figure 4b**). (Co-administration of the 5 mg/kg dose of K1592 with MK-801 was not performed due to side effects.)

Regarding K1594, the analysis revealed the effect of K1594 and MK-801 (F (2, 44) = 15.54 and F (1, 44) = 68.95, respectively, p < 0.0001 for both). At the dose of 5 mg/kg, K1594 decreased locomotion of the intact as well as MK-801-treated rats (p = 0.0074 and p < 0.0001, respectively), while at the lower dose it did not affect locomotion of any of these groups (**Figure 4c**).

Finally, in the K1599 group, the analysis showed the effect of K1599 (F (2, 44) = 4.663, p = 0.0146) and MK-801 (F (1, 44) = 115.4, p < 0.0001). At the dose of 5 mg/kg, K1599 mitigated the MK-801-induced hyperlocomotion (p = 0.0044 vs. VEH + MK-801), while not affecting locomotion of intact animals. At the lower dose, it had no effect on locomotion in any group (**Figure 4d**).

### 3.3 *In situ* acetylcholinesterase activity assay

K1578, K1592, K1594, and K1599 are AChE inhibitors displaying IC_50_ values of 1.58, 0.223, 0.072 and 8.22 µM, respectively, as proven by a previous *in vitro* assay utilizing modified Ellmann’s method and human recombinant AChE ^27^. To investigate mechanisms possibly underlying the results from the scopolamine-induced animal model, the effects of the compounds (1 mg/kg ip) on the *in situ* AChE activity in selected rat brain structures were assessed.

The spectrophotometric analyses of AChE activity did not reveal any significant alterations in either the hippocampus (**Figure 5a**) or the prefrontal cortex (**Figure 5b**) upon treatment with the studied compounds.

**Figure 5.**
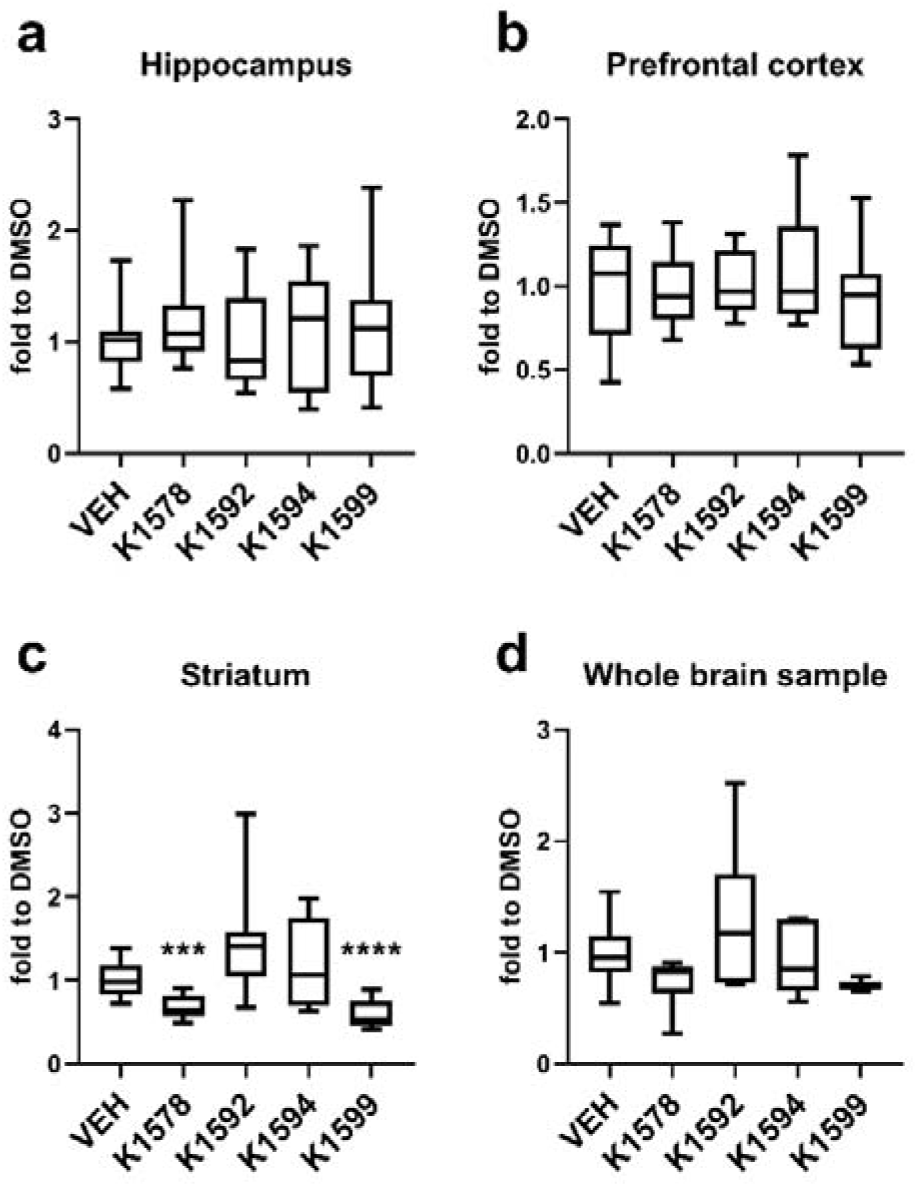
Acetylcholinesterase activity. The effect of the compounds (1 mg/kg ip) on AChE activity in the hippocampus (**a**), prefrontal cortex (**b**), striatum (**c**) and whole brain sample (**d**). K1578 and K1599 decreased the AChE activity in the striatum. VEH – vehicle. Data are represented as the median with minimum to maximum range, * vs. VEH, *** p < 0.001, **** p < 0.0001. VEH samples AChE enzyme activities reached the following absolute values (a) 15.19 ± 3.09 U/mg protein, (b) 9.810 ± 1.54 U/mg protein, (c) 26.05 ± 3:27 U/mg protein and (d) 27.38 ± 3.36 U/mg protein.

In stark contrast, the analyses unveiled a substantial reduction in AChE activity within the striatum (W (4.000, 18.85) = 12.77, p < 0.0001). Specifically, in the case of K1578, AChE activity decreased by 32% (p = 0.0003), and for K1599, it decreased by 42% (p < 0.0001). The results hence suggest that the selected dose levels of these compounds induce brain region-specific alterations in AChE activity. Conversely, neither K1592 nor K1594 elicited any statistically significant changes in AChE enzyme activity within the striatum when compared to the control group (**Figure 5c**).

Regarding the whole brain sample, the analysis revealed the treatment effect (F (4, 38) = 3.741, p = 0.0116), but the Dunnett’s multiple comparisons test failed to find differences between the groups (**Figure 5d**).

### 3.4 *In vitro* electrophysiology

K1578, K1592, K1594, and K1599 act as NMDA receptor inhibitors. At membrane potential of –60 mV they possess IC_50_ values of 21.01, 7.29, 17.05 and 4.16 µM, respectively, for GluN1/GluN2A receptors, and 8.69, 22.07, 7.83 and 14.56 µM, respectively, for GluN1/GluN2B receptors ^27^. As only K1599 ameliorated the MK-801-induced pathologies in the current study, a more detailed electrophysiology study focusing on the interaction of K1599 with GluN1/GluN2A receptors was conducted, in order to further investigate the mechanism possibly underlying its beneficial effects.

Whole-cell patch-clamp measurements on transfected HEK293 cells expressing rat version of the GluN1-4a/GluN2A (GluN1/GluN2A) at different membrane potentials showed that K1599 inhibits the GluN1/GluN2A receptors at all studied membrane potentials. However, the inhibitory effect was more pronounced at the negative membrane potentials (**Figure 6a,b**).

**Figure 6.**
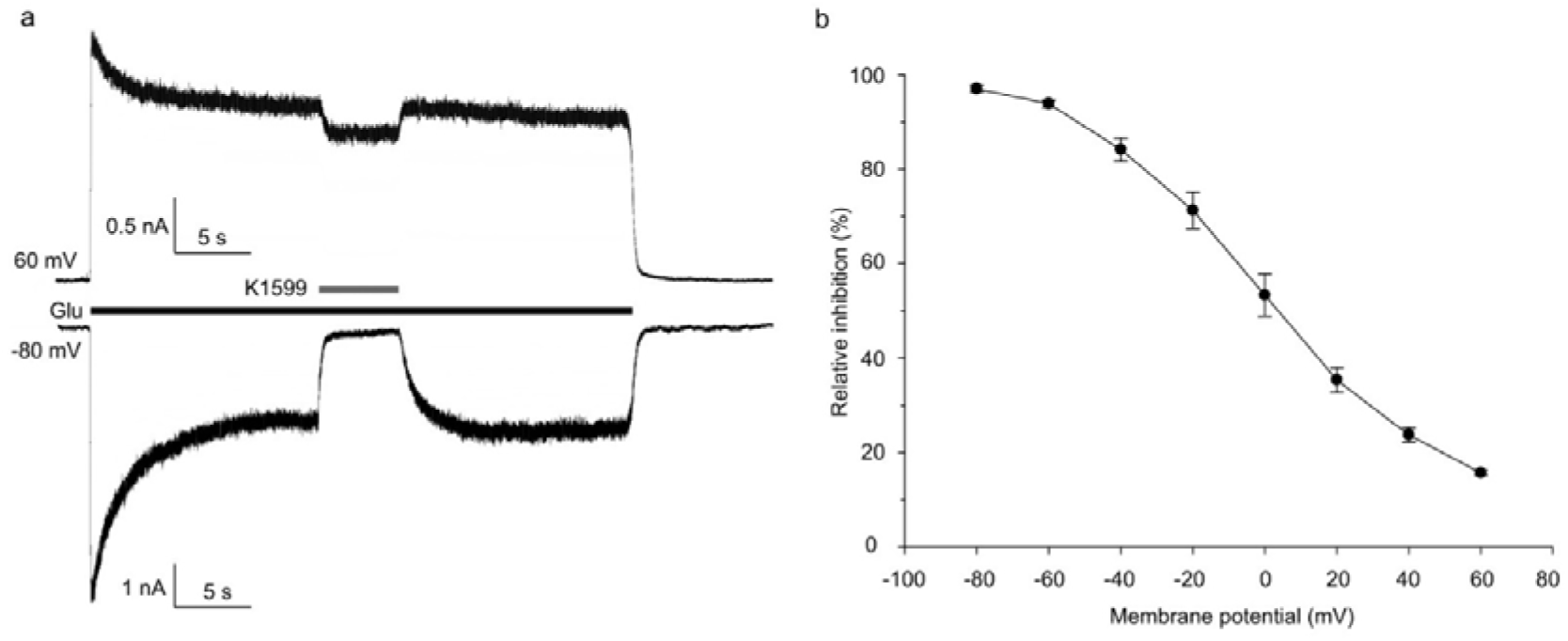
Electrophysiology: The inhibition of GluN1/GluN2A receptors by K1599. Representative whole-cell patch-clamp recordings measured from HEK293 cells expressing the GluN1/GluN2A receptors held at a membrane voltage of −80 mV and +60 mV; 30 μM K1599 was applied as indicated (**a**). Graph summarizing the relative inhibition induced by 30 µM K1599, measured at the indicated membrane potentials. *n* ≥ 5 cells per each condition (**b**).

## 4 Discussion

The current study revealed different *in vivo* effects of acute pretreatment with tacrine derivatives acting as dual inhibitors of cholinesterases and NMDA receptors ^27^ in cholinergic and glutamatergic animal models of cognitive deficit.

### 4.1 Cholinergic effects

The first part of the study focused on the interactions of the compounds with the cholinergic system as acetylcholine modulates learning and memory processes ^52,53^. Consistently with literature, scopolamine induced cognitive deficit of both spatial learning in acquisition trials ^35,39,54,55^ and reversal learning ^55,56^ in the MWM. It is known that these deficits can be mitigated by indirect stimulation of the cholinergic system by various AChE inhibitors including tacrine ^35,39,54,55^. In the current study, K1578 and K1599 alleviated the reversal learning deficits, with K1599 being active over a wider dose range (both tested doses). On the other hand, none of the compounds was effective against the deficit of spatial learning in acquisition trials.

To understand these effects, an *in situ* AChE inhibition study was conducted in separate sets of animals administered with the compounds at 1 mg/kg via the same route and timing as the rats subjected to MWM. We focused primarily on hippocampus, important for spatial learning during MWM acquisition trials ^46,57^ and striatum, mediating spatial reversal learning and cognitive flexibility in general ^45,58,59^. In particular, intact cholinergic system in these structures seems to be crucial, as proven by the disrupting effect of locally (intrahippocampally or intrastriatally) infused scopolamine on spatial discrimination learning ^60^ and reversal learning ^61^, respectively.

In the hippocampus none of the compounds inhibited AChE significantly, possibly explaining the lack of impact on spatial learning. It is unclear whether the compounds are fully ineffective, or different doses than those used in our study would be necessary to restore spatial memory. The known inverted U-shaped dose-response relationship and narrow therapeutic window of many AChE inhibitors ^35,62–64^ raise the possibility that the latter may be true. Similarly as in the hippocampus, AChE activity in the prefrontal cortex remained unchanged.

However, K1578 and K1599 significantly inhibited AChE (by 32% and 42%, respectively) in the striatum. The same dose of these compounds also mitigated the scopolamine-induced deficit of reversal learning in the rats subjected to the MWM. There is strong evidence that the cholinergic system in the striatum plays a key role in reversal learning, but not in initial acquisition ^45,59,61^. Specifically, cholinergic interneurons in dorsomedial striatum are assumed to be crucial for establishment of new strategies when conditions change ^45^. Therefore, we suggest that there can be a causal relationship between inhibition of striatal AChE by K1578 and K1599 and their positive effects on reversal learning.

Our results hence indicate that K1578 and K1599 induce different degrees of AChE inhibition in different brain structures. Interestingly, a similar structure-dependent effect was observed with tacrine. Systemic administration of tacrine induced a relatively higher level of AChE inhibition in striatum than in hippocampus, while the opposite was true for another AChE inhibitor rivastigmine ^35^. Correspondingly, lower dose of tacrine is sufficient to increase acetylcholine level in striatum, than in the hippocampus or cortex ^65^. Thus, brain structure-dependent effects of AChE inhibitors may lead to their different effects in individual cognitive tasks.

Whole brain AChE inhibition was not significant for any compound, suggesting the importance of individual brain structures. Unexpectedly, K1592 and K1594, despite lower *in vitro* IC_50_ values for (human recombinant) AChE than those of K1578 and K1599 ^27^, did not inhibit AChE in any brain structure. This discrepancy may perhaps relate to differences in distribution to the brain ^27^, emphasizing the importance of *in situ* AChE activity assessment.

In summary, K1578 and K1599 exhibit brain structure-specific AChE inhibition, which may explain their positive effects on reversal learning in the scopolamine-induced model. Our findings also highlight the need for *in situ* AChE activity assessment in preclinical research in parallel with the behavior experiments.

### 4.2 Glutamatergic effects

The second part of the study focused on the interactions of the compounds with the glutamatergic system, specifically NMDA receptors. Consistently with literature, MK-801, a potent NMDA receptor antagonist, impaired MWM acquisition ^41,66^, underlining that intact NMDA receptors are crucial for learning and memory ^67,68^. However, the selected dose of MK-801 did not impair reversal learning. Literature dealing with the effects of MK-801 on spatial reversal provides conflicting results ^69–71^. Of the compounds studied, K1599 (1 mg/kg) mitigated the MK-801-induced deficit of spatial learning, possibly by competing with MK-801 on NMDA receptors.

NMDA receptor antagonists represent a heterogeneous group of compounds with diverse biological effects. The NMDA receptors are essential for physiological functions including memory, but their over-activation is implicated in neuropathological processes ^10^. Therefore NMDA receptor antagonists may possess therapeutic, but also side effects, depending on precise mechanisms of interaction with the receptor ^72–75^.

High-affinity open-channel blockers like MK-801 interfere with physiological function of NMDA receptors and hence impair memory and induce other side effects like hyperlocomotion ^41,76,77^. On the other hand, some open-channel blockers with moderate affinity, like memantine and 7-MEOTA, or some NMDA receptor subunit-selective antagonists, may possess lower risk of side effects ^25,26,72–74,78,79^, and even positive effects on cognition ^10,77,80^. Our tacrine derivatives previously showed no side effects typical of NMDA receptor antagonists ^27^. When co-administered, the NMDA receptor antagonists can interact complexly, either potentiating or mitigating behavioral effects of each other ^73,78^. Counterintuitively, some NMDA receptor antagonists can reverse the MK-801-induced cognitive deficits ^74,81^.

In our study, K1599, uniquely effective against the MK-801-induced cognitive deficit, may owe its efficacy to brain accumulation and superior brain availability compared to the other studied compounds ^27^. Moreover, its NMDA receptor subunit-dependent action, namely preferential inhibition of GluN1/GluN2A over GluN1/GluN2B receptors, ^27^, could also contribute.

Most NMDA receptors are heterotetramers containing GluN1 and GluN2A–D subunits. GluN2A and GluN2B represent the most abundant GluN2 subunits in the cognition-related structures in the adult brain ^82,83^. However, they show different functional properties and roles ^83^ and their selective antagonists may exert different behavioral effects ^77^. Regarding our study, MK-801 potently inhibits GluN1/GluN2A and GluN1/GluN2B receptors ^84^, while our tacrine derivatives are less potent and show slight preference for GluN1/GluN2A (K1592, K1599) or GluN1/GluN2B (K1578, K1594) receptors ^27^. GluN2B-selective antagonists seem potentially appropriate as AD drugs, as these receptors are supposed to play role in its pathophysiology ^83^ and contribute more intensely to the neuronal injury than GluN2A-containing receptors ^28,29^. In view of that, the pro-cognitive effect of the GluN1/GluN2A-preferring antagonist K1599 may seem surprising. On the other hand, long term potentiation and hence presumably also memory processes are dependent predominantly on GluN2A-(rather than GluN2B-) containing receptors ^85,86^. Therefore, we hypothesize that preferential competition of K1599 with MK-801 for binding on GluN2A-containing receptors may perhaps contribute for mitigation of cognitive deficit. Nevertheless, it is not clear whether the GluN2A-preference of K1599 is indeed responsible for its pro-cognitive effect.

Next, the effects of the compounds on locomotion of intact animals and animals with MK-801-induced hyperlocomotion ^72,78^ were studied in the open field. Compounds decreasing intact animals’ locomotion (K1578 5 mg/kg, K1592 1 mg/kg, K1594 5 mg/kg) also mitigated the MK-801-induced hyperlocomotion, suggesting a non-specific effect, not necessarily mediated via NMDA receptors. AChE inhibitors can induce similar effects, as known from literature ^36,87,88^ as well as it was observed in our laboratory with the AChE inhibitor donepezil (unpublished data). Therefore, the AChE-inhibitory effect of the compounds may be perhaps responsible.

K1578 (1 mg/kg) and K1599 (5 mg/kg) displayed interesting interactions with MK-801, potentially reflecting their subunit-dependent NMDA receptor inhibition. GluN2B-preferring compound K1578 slightly potentiated the MK-801-induced hyperlocomotion, at a dose, which did not affect locomotion of intact animals (1 mg/kg). Interestingly, a similar effect was described with another GluN2B-preferring NMDA receptor antagonist ifenprodil ^78^. On the other hand, K1599 mitigated the MK-801-induced hyperlocomotion at a dose, which did not affect locomotion of intact animals (5 mg/kg). Similar effect was described with 7-MEOTA and interpreted as competition of 7-MEOTA and MK-801 for the binding site on NMDA receptors ^25^. The effect of K1599 and 7-MEOTA may be perhaps related to their GluN2A-preference ^25,27^, as the locomotion-stimulating effect of MK-801 seems to be mediated greatly via inhibition of GluN2A-containing NMDA receptors ^84^. The open field and MWM results seem to be consistent with the notion that K1599 affects NMDA receptors in a way that does not impede their physiological function, but by competition with MK-801 for binding at NMDA receptors it mitigates the detrimental effects of MK-801.

The findings from our *in vitro* electrophysiology study align with this concept. The results indicated that K1599 inhibits the GluN1/GluN2A receptors at all studied membrane potentials, with a more pronounced effect at the negative membrane potentials. This conclusion is in agreement with our previous data ^27^. While it is technically challenging to directly show the competition between the K1599 and MK-801 using *in vitro* electrophysiology because of the “irreversible” nature of the MK-801 block ^89^, we conclude that both K1599 and MK-801 likely act as open-channel blockers, and thus, they could compete for the same binding site within the ion channel region of the GluN1/GluN2A receptor *in vivo*.

### 4.3 The dual effect of K1599: From structural modifications back to 7-MEOTA-like compounds

To sum up, K1599 demonstrated the most favorable effects among the compounds studied, mitigating cognitive deficits induced by disruptions to both the cholinergic and glutamatergic systems at a fixed dose (1 mg/kg). This aligns with its intended dual cholinesterase- and NMDA receptor-inhibitory effect.

Interestingly, of the compounds studied, the favorable compound K1599 possesses the most similar structure and affinities for target proteins as 7-MEOTA. K1599 differs from 7-MEOTA only by the smaller size of the cycloalkyl moiety attached to the aromatic region. 7-MEOTA ^22,25^ and K1599, in contrast with the other compounds from the current study, showed beneficial *in vivo* effects in the glutamatergic model, consistent with NMDA receptor inhibition. K1599 and 7-MEOTA display balanced IC_50_ values for AChE, BChE and GluN1/GluN2A and GluN1/GluN2B receptors (at – 60 mV), differing by no more than one order of magnitude ^25,27^, which is considered one of the fundamental feature of multi-target drugs ^12,14^. The IC_50_ values of K1599 display following relationship: GluN1/GluN2A < AChE < BChE < GluN1/GluN2B (4.16; 8.22; 10.6; 14.56 µM, respectively, with inhibition of NMDA receptors measured at the membrane potential of – 60 mV) ^27^. Both K1599 and 7-MEOTA show slight preference for GluN1/GluN2A over GluN1/GluN2B receptors ^25,27^. Besides, the higher brain availability of K1599 compared to the other compounds from the current study ^27^ may contribute to its superior effects when given in equal doses.

On the other hand, the compounds K1578, K1592, and K1594 were not beneficial. They differ from K1599 by different substituents at the core moiety, and the size of the cycloalkyl moiety. Contrary to K1599, these compounds display preferential inhibition for cholinesterases over NMDA receptors. K1578, with chlorine atom substitution in the position identical to methoxy substitution in 7-MEOTA, was effective only in the cholinergic model. K1592 and K1594, displaying increased inhibitory potency for AChE or both AChE and BChE *in vitro* ^27^, were not considerably effective in either model. To sum up, our results indicate that tacrine derivatives with structural and functional similarity to 7-MEOTA may be especially promising in the development of AD drugs with dual *in vivo* effects.

### 4.4 Study limitations

The study has several limitations. Firstly, the use of the MK-801-induced model lacks a direct link to AD; instead, it served as a pharmacological tool to explore NMDA receptor-mediated behavioral effects. Consequently, translating our findings, such as the role of NMDA receptor subtype preference of the compounds, to AD patients remains uncertain.

Secondly, although the scopolamine- and MK-801-induced models were employed to investigate AChE- and NMDA receptor-mediated effects, respectively, it is essential to recognize the close interaction between both neurotransmitter systems (see ^10^). It was reported that in some cases, cholinergic agents can indirectly affect the glutamatergic system ^36,88,90^, while glutamatergic agents can influence the cholinergic system ^16,91–93^. Tacrine derivatives’ effects in both models might hence result from intricate cholinergic and glutamatergic interactions, and involvement of other neurotransmitter systems cannot be ruled out.

Lastly, our compounds vary not only in affinities for target proteins but also in brain availability, potentially influencing behavioral outcomes. This challenges a direct assessment of the structure-activity relationship. While intracerebroventricular administration could address this issue, it is deemed inappropriate for the extensive screening study due to invasiveness, clinical irrelevance, and time demands.

## 5 Conclusions

Our comprehensive *in vivo* study delved into the effects of tacrine derivatives, functioning both as cholinesterase inhibitors and NMDA receptor antagonists, utilizing animal models reflective of cognitive deficits arising from cholinergic or glutamatergic dysfunction. K1599 emerged as the most promising compound, demonstrating efficacy at a consistent dose across both models, hence confirming its dual *in vivo* effect. Deciphering the structural and functional attributes of tacrine derivatives associated with optimal *in vivo* pro-cognitive effects holds potential for advancing the development of dual compounds as promising therapeutics for AD.

## Abbreviations

AChE: acetylcholinesterase
AD: Alzheimer’s disease
ANOVA: analysis of variance
BChE: butyrylcholinesterase
DMSO: dimethyl sulfoxide
DTNB: 5,5′-dithiobis-2-nitrobenzoic acid
IC_50_: the half maximal inhibitory concentration
K1578: 7-chloro-1*H*,2*H*,3*H*-cyclopenta[b]quinolin-9-amine hydrochloride
K1592: 1-chloro-6*H*,7*H*,8*H*,9*H*,10*H*-cyclohepta[b]quinolin-11-amine hydrochloride
K1594: 6-methyl-1,2,3,4-tetrahydroacridin-9-amine hydrochloride
K1599: 7-methoxy-1*H*,2*H*,3*H*-cyclopenta[b]quinolin-9-amine hydrochloride
MK-801: dizocilpine
MWM: Morris water maze
NMDA receptor: N-methyl-D-aspartate receptor
SEM: standard error of the mean
VEH: vehicle-treated group
7-MEOTA: 7-methoxytacrine

## Authors’ contributions

MC: Writing - original draft; Writing - review & editing, Formal analysis, Investigation - behavioral study, Conceptualization. DK: Writing - review & editing, Investigation - biochemistry, Formal analysis. KK: Investigation - biochemistry. LC: Investigation - behavioral study. AM: Investigation - electrophysiology. KH: Investigation - behavioral study. LG: Resources. MH: Funding acquisition. JK: Resources, Funding Acquisition. OS: Funding acquisition. KV: Funding acquisition, Conceptualization, Supervision, Writing - review & editing.

